# A low-cost method for collecting and measuring sinking speed of particles in coastal zones

**DOI:** 10.64898/2026.06.11.730824

**Authors:** Josephin Lemke, Kristian Spilling

## Abstract

Sinking marine particles is a key process regulating carbon export through the biological carbon pump, yet direct measurements of sinking dynamics remain limited in many coastal environments. One barrier is that most existing approaches require expensive instrumentation and large research platforms. Here, we present a low-cost, modular method for concentrating fast-sinking particles and measuring their individual sinking velocities under controlled conditions. This combines large settling tanks (110 L) for field-based particle fractionation with a video-based tracking system that quantifies the sinking behavior of natural marine particles. The particle sinking speed chamber is surrounded on three sides by a temperature-controlled water chamber, minimizing the problem of advection during measurements. The post-processing Python script delivers sinking velocity, particle size, circularity, and RGB-based properties for large numbers of particles. The method accuracy was validated using reference beads with known theoretical sinking velocities derived from Stokes’ law. Field deployments in the Baltic Sea demonstrated successful enrichment of fast-sinking particles and stable operation from both a research vessel and a small boat. Compared to existing methods, the approach substantially reduces logistical and financial barriers while maintaining particle-resolved measurements and compatibility with complementary biogeochemical analyses. This enables a broader observational coverage of sinking particle processes across environments that are currently underrepresented in carbon export studies.

## Introduction

Marine particles and aggregates form the biological carbon pump (BCP), which is essential in the removal of carbon dioxide (CO_2_) from the atmosphere (Fowler and Knauer 1986). As part of the BCP, phytoplankton fix dissolved inorganic carbon into organic matter via photosynthesis. This organic matter forms the basis for sinking particles. These particles can aggregate into marine snow and transport carbon from surface waters to deeper layers. In this way, they contribute to long-term carbon sequestration and the reduction of atmospheric CO₂ concentrations (e.g., Volk and Hoffert 1985; Ducklow et al. 2001).

Marine aggregates provide substrates for heterotrophic microorganisms and zooplankton to feed on, resulting in remineralization of organic matter, releasing the carbon as CO_2_ back into the water column during the respiration process (Simon et al. 2002; Steinberg et al. 2008). The efficiency of the BCP is mainly determined by the balance between remineralization and sinking speed. Physical forcing such as upwelling or downwelling can additionally influence particle transport pathways. The fate of aggregates also depends on their location in the water column. In the mixed layer, microbial activity and turbulence allow for transformation and remineralization, whereas below the mixed layer conditions may allow a larger fraction of OM to be transported downward without rapid degradation (e.g., Heiskanen 1998; Francois et al. 2002; Stocker 2012).

The fate of marine particles is controlled by their physical properties. Aggregate size, density, shape, porosity, and composition determine the sinking speed and thus influence how long they remain in the water column (e.g., Laurenceau-Cornec et al. 2015; Iversen and Lampitt 2020; Omand et al. 2020). In addition, processes such as aggregation, disaggregation, ballasting, and microbial degradation continuously modify particle structure, sinking velocities, and lifetime (Briggs et al. 2020). The physical properties of the sinking particles, such as porosity, could affect both remineralization and sinking speed to contrasting directions (Spilling et al. 2023), and, sinking velocity is not constant but changes over time and across environments. This makes sinking speed a key parameter for understanding how efficiently organic carbon is exported to depth (Chajwa et al. 2024).

This role of sinking velocity and its influence on the BCP becomes particularly important in coastal environments, which are considered a net sink for atmospheric CO_2_ (Bauer et al. 2013). These areas are characterized by high primary productivity, which increases the availability of organic matter and promotes aggregation process that can enhance rapid particle export. In addition, riverine discharge supplies substantial amounts of particulate organic carbon to coastal waters (Bauer et al. 2013). Coastal environments are also characterized by elevated inputs of mineral particles, which can enhance carbon export efficiency by acting as ballast and increasing particle density and sinking velocity (Iversen and Ploug 2010; Rixen et al. 2019; Mueller et al. 2022). In combination with shallow depths, this increases the probability of carbon reaching the seafloor before being remineralized, as particle flux generally decreases with depth during sinking according to the Martins curve (Martin et al. 1987; Buesseler and Boyd 2009). Parts of the deposited organic matter will be remineralized within the sediments, but parts can be permanently buried. Coastal systems act as important sites for carbon sequestration despite covering a relatively small area (∼8%) of the global ocean (Longhurst 1995; Gattuso et al. 1998; Bourgeois et al. 2016), while they account for roughly one tenth of the present-day global ocean CO₂ sink (Dai et al. 2022). These processes (particle sinking, remineralization, and burial) are central to the functioning of the BCP and determine the efficiency of long-term carbon sequestration.

Coastal environments are highly dynamic characterized by strong spatial and temporal variability in carbon cycling and fluxes, and the interplay of multiple physical and biogeochemical processes (Bauer et al. 2013). Resuspension, lateral transport, turbulent mixing, and biologically mediated transports complicate the pathways and fate of sinking particles (e.g., Blair and Aller 2012; Bauer et al. 2013; Nooteboom et al. 2020). This complexity makes it difficult to resolve carbon fluxes and particle transport, highlighting the need for accessible observational approaches that can be applied across diverse coastal environments and over extended spatial and temporal scales.

To quantify sinking velocities of marine particles, several methods have been developed. In situ methods, such as camera systems and holographic imaging, allow direct observation of particles in their natural environment and high ecological realism (e.g., Graham and Nimmo Smith 2010; Simoncelli et al. 2019; Soviadan et al. 2025). However, these methods often do not provide information on the composition of particles. Indirect approaches, such as sediment traps, offer detailed information on biogeochemistry but not on the sinking of individual marine particles (McDonnell and Buesseler 2010; Laurenceau-Cornec et al. 2015). Laboratory-based methods, in which sinking velocities are measured ex situ, such as sedimentation columns (Bach et al. 2012; Laurenceau-Cornec et al. 2015) or vertical flow systems (Ploug et al. 2010; Flintrop et al. 2024), offer controlled measurements coupled with biogeochemical analysis. However, these methods may alter the natural particle properties and natural sinking behavior due to handling, simplified hydrodynamic conditions, and artificial water movements withing the measurement system. As a result, the approaches to quantify sinking velocities often involve compromises between realism and experimental control.

Beyond these methodological trade-offs, practical constraints further limit the application of existing approaches. Many methods rely on specialized instruments, large research vessels, or laboratory facilities that are not always available outside well-funded research projects. This is especially limiting in coastal areas, where fieldwork is often carried out on smaller platforms and with limited resources. As a result, direct measurements of marine particle sinking speed are relatively rare in these settings. There is therefore a need for simpler and more accessible approaches that can be applied in the field without complex infrastructure, while still providing reliable and reproducible results.

In this study, we present a system for measuring sinking velocities of natural particle assemblages under controlled conditions. The method includes field-based sampling, where suspended and fast-sinking particles are separated in a settling tank, and laboratory-based measurements of individual particle sinking behavior using optical tracking. Laboratory measurements allow us to relate sinking velocity to particle-specific properties such as size, shape, while also extracting RGB-based particle information. In addition, bulk analyses of suspended and fast-sinking fractions provide complementary information on their overall biogeochemical composition. We used low-cost components off the shelf from hardware and electronic shops. By reducing technical and financial barriers, we aim to make carbon export measurements more accessible across a wider range of environments, especially targeting coastal environments that remain underrepresented in current observational datasets.

## Materials and procedures

### Particle fractionation in settling tank

To concentrate the fast-sinking particles, we constructed two 110 L settling tanks (CIPAX, Sweden, Figure 1). Each tank was equipped with taps at varying heights, enabling water withdrawal from different levels within the tanks, including one tap at the bottom. A custom-built, wooden frame was used to keep the tanks stable and use the tap underneath the tanks.

**Figure 1:**
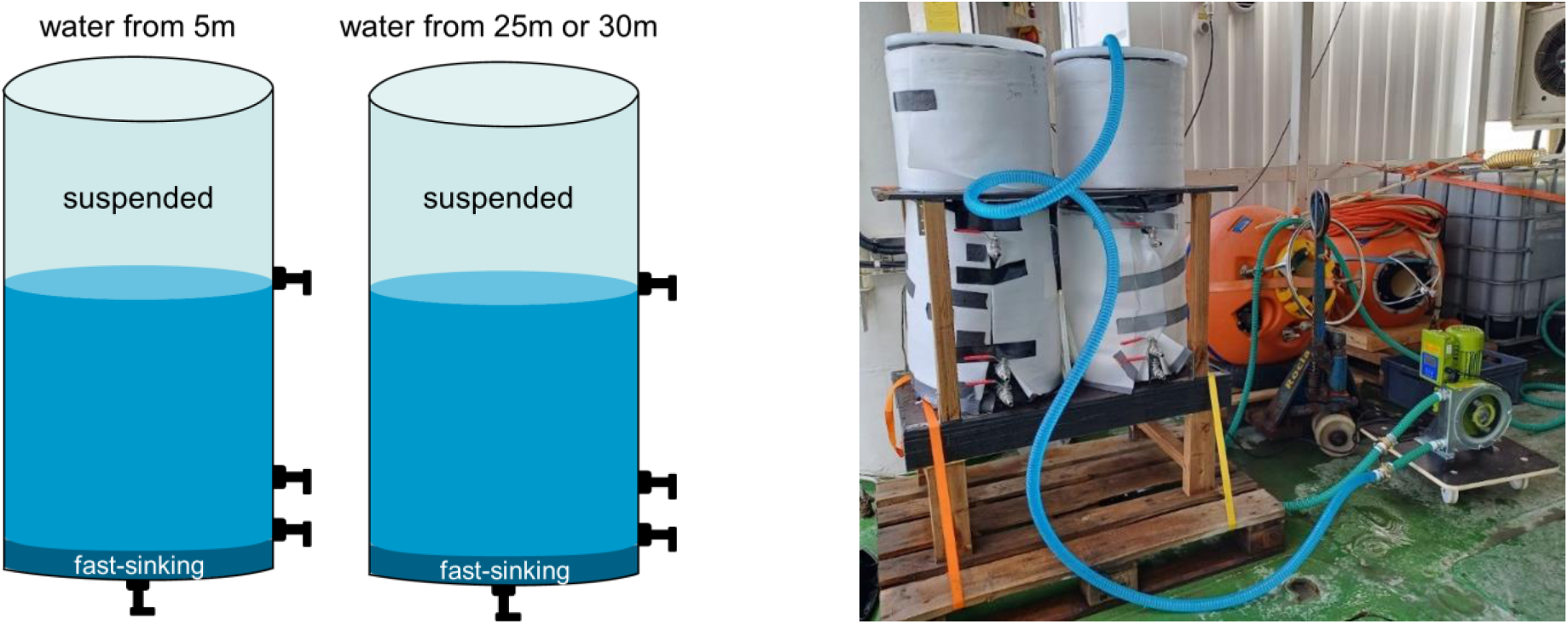
The settling tanks schematics (left) and how they were mounted inside a wooden holding frame (right). The tanks were covered with optical and thermal insulation and were filled with water from 5 m and 30 m depth with a peristaltic pump. They feature multiple taps at varying heights, allowing for sampling from the suspended fraction and fast-sinking fraction.

During trial testing, the tanks were filled with seawater collected from two depths, 5m and 30 m. Water from 5 m was chosen to represent the surface mixed layer. Water from 30 m depth was collected to represent water typically below the mixed layer in our Baltic Sea test environment. The settling tanks were gently filled with a peristaltic pump system (Albin ALP17N pump, France), that transferred water directly from depth into them via PVC spiral tubing (APD, with inner diameter Ø = 2.5 cm). The tanks were filled completely, and the lids were strapped on, leaving no head space. After filling, the tanks were left undisturbed for two hours to allow particles to settle (Riley et al. 2012). To minimize photosynthesis and temperature effects, the tanks were shielded from light with an optical insulating layer (black shield sheet) and covered with thermal insulation material against heat exposure.

Following the two-hour settling period, particle fractions were sampled sequentially. Water collected from the upper tap, representing the upper half of the tanks (8 L), was defined as the suspended fraction. Two intermediate taps were opened during sampling, but this water was discarded. Water collected from the bottom tap (∼5.5 L) represented the fast-sinking fraction, which was enriched in rapidly settling particles. All samples were transferred into four 8 L Nalgene bottles. As the focus of the subsequent analyses was on sinking material, only the fast-sinking fraction was used.

### Set up of sinking speed measurements

The particle sinking speed chamber (PSSC; Figure 2) used to measure sinking velocities was made from polycarbonate roofing boards (Gutta Werke, Easy Click, Germany). Each board consists of 12 parallel channels (16 x 16 mm), from which sections of 40 cm length were cut. One central channel was used as the PSSC. To enable temperature control, two boards were glued together (2-component epoxy glue) after removing the walls of the channels that surrounded the central settling channel, and all seams were closed with transparent silicone sealant. This created a connected chamber covering three sides of the PSSC that was used for temperature control (TC chamber). This TC chamber was connected to a circulating water temperature control unit (LAUDA RE 420 SN; Supporting Information Fig. S1), with inlet and outlet ports (general purpose tube connectors glued in place) located at the top and bottom of the TC chamber, respectively. That allows continuous flow and stable in situ temperature conditions during measurements. The connected boards were mounted on a custom build frame made from 4 x 4 cm wooden boards.

**Figure 2:**
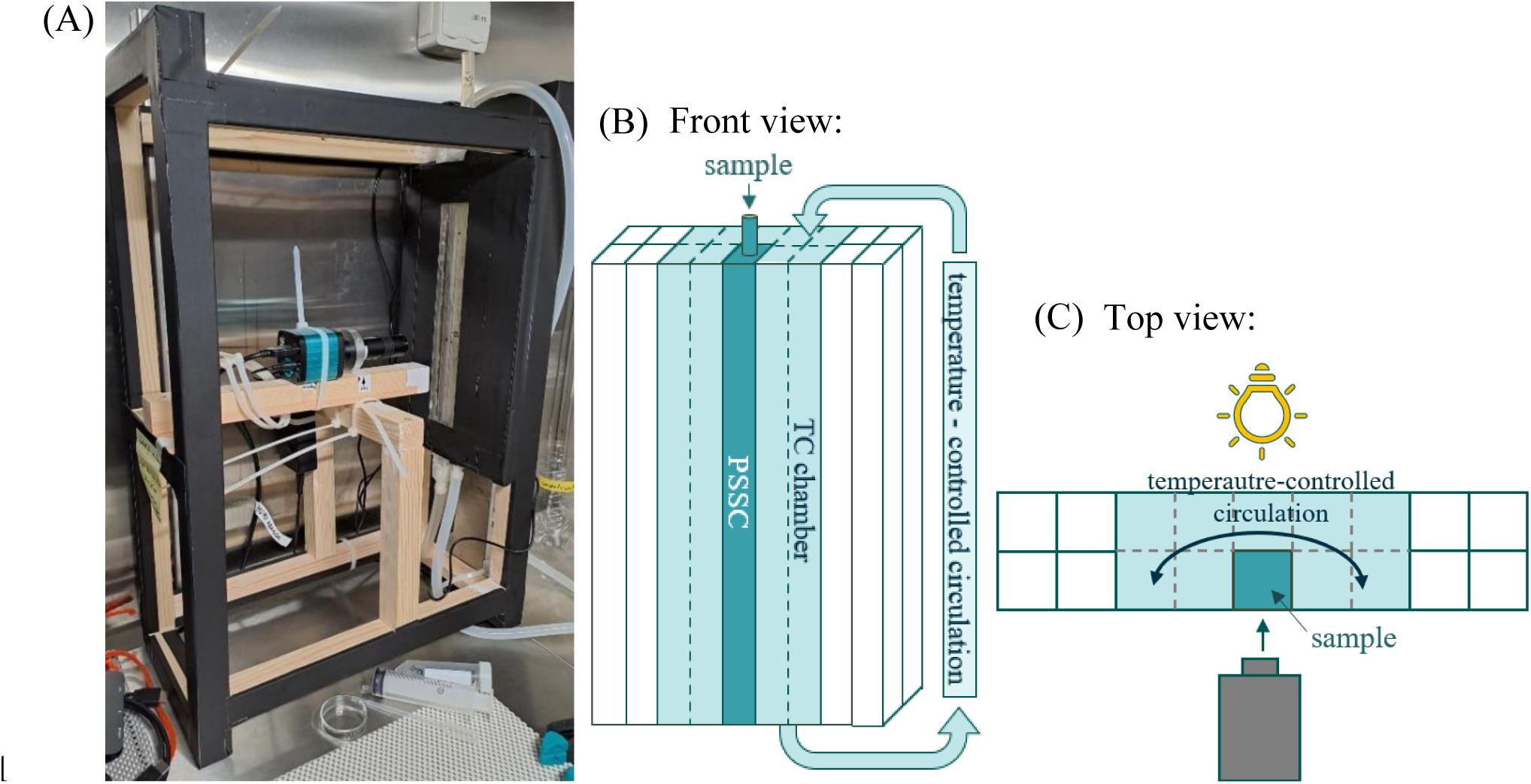
Photograph and schematic figure of the experimental setup used for sinking velocity measurements. (A) Board with particle sinking speed chamber (PSSC) and temperature control chamber (TC chamber) mounted on the supporting wooden frame, in front of the camera. (B) Schematic front view of the boards, including the central PSSC, the sample inlet at the top, and the surrounding temperature control circulation. Dashed lines indicate the removed channel walls forming the temperature control compartment. (C) Schematic top view, showing the position of the light source behind the PSSC, the camera in front, and the temperature control circulation surrounding the PSSC.

Prior to measurements, the PSSC was carefully filled with sampled water from the top while avoiding air bubbles. The inlet hose was then sealed with a clamp to prevent air from entering the system. This ensured that the system was fully filled with water. A hose connected to the bottom of the PSSC allowed controlled drainage and rinsing between measurements. A camera (HIGH CLOUD, China) was mounted directly in front of the settling channel to record particle movement, while a light source (integrated camera USB LED light) placed behind the panel illuminated the particles. A microscale placed in front of the PSSC was used to calibrate the camera and determine the pixel dimension before the camera was moved forward to keep the focus in the middle of the PSSC. The recorded videos were subsequently used to track individual particles and determine their sinking velocities.

### Video processing and particle tracking

The recorded videos were analyzed using a Python-based image processing workflow to track individual particles and quantify their sinking behavior. The analysis consisted of the detection of single particles, the reconstruction of the trajectories, and the subsequent post-processing of particle velocities and properties. The complete workflow is provided as Script S1 in the Supporting Information.

#### Particle detection and tracking

Particles in each frame were identified using a background subtraction approach based on a Gaussian mixture model (OpenCV library). The method models the intensity distribution of each pixel over time and detects deviations from the background as moving particles by marking extreme value pixels. These results are then denoised using morphological filtering to remove small, weakly connected detections. This step establishes an effective lower limit on particle size, rather than a fixed size threshold. Detected particles are tracked across consecutive frames by linking each particle to the nearest detection in the subsequent frame within a defined search radius (20 pixels). Tracking was performed on a one-to-one basis. Therefore, particle merging and splitting events were not resolved. When particles merged, the path of one trajectory was stopped, and when particles split, a new trajectory was initialized. Each tracked particle was assigned a confidence score that increased when the particle was successfully detected in consecutive frames and decreased when it was not detected. Tracking was stopped when the confidence score reached zero. Only particles that were for a user defined minimum duration were retained for further analysis to exclude short-lived or unreliable detections.

#### Trajectory analysis and velocity calculation

For each tracked particle, the position of its center (derived from the bounding box) was recorded throughout its entire detection. Sinking velocity was calculated from the net vertical displacement between the first and last detected particle position. This approach avoids overestimating sinking velocities, as lateral particle motion increases the distance travelled without contributing to vertical sinking. The displacement in pixels was converted to physical distance using a calibration factor of 720 pixels corresponding to 5.8 mm (0.00806 mm px⁻¹, C_px_). Particle lifetime was defined as the number of frames over which a particle was tracked and converted to time using the frame rate of the recordings (frames per second, fps). Sinking velocities were thus calculated as:

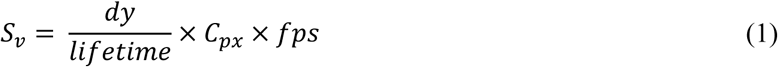

The resulting velocities were converted from mm s^-1^ to m day⁻¹.

#### Post-processing

Particle sinking velocities decrease close to chamber walls because of increased hydronamic drag (Happel and Brenner 1983; Uhlherr and Chhabra 1995). Velocities were therefore corrected for wall effects following (Ristow 1997):

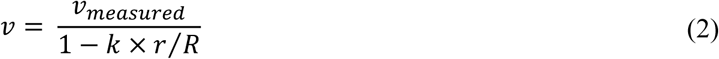

where v is the corrected sinking velocity, v_measured_ is the measured sinking velocity, k the drag coefficient, r the particle radius, and R the distance between particle and chamber wall. Following Brenner 1962 and Bach et al. 2012, the drag coefficient was assumed to be k = 1.004 for particles sinking approximately midway between two parallel walls. The focus of the camera was set to the center of the chamber, where the distance to the wall was R = 0.875 cm. This correction equation has previously been shown to improve the consistency between measured and theoretical sinking velocities of reference particles (Bach et al. 2012). However, the same study showed that the resulting wall-effect corrections were generally small, although they increased with particle size.

To identify sinking particles and distinguish them from particles with pronounced lateral movement, trajectories were classified based on their net displacement during the tracking period. Particles were considered sinking when their final vertical position was lower than their initial position and their horizontal displacement remained 207 µm. This filtering step excluded particles with substantial sideways movement and retained particles showing predominantly vertical sinking. Only particles meeting these criteria were included in subsequent analyses of sinking velocity and particle properties.

Additional particle properties were calculated from the particle contours identified during the image processing. Particle size was characterized using the area-based diameter (ABD) and equivalent spherical diameter (ESD). ESD was calculated as the mean Feret diameter based on measurements taken at 36 orientations. Particle shape was characterized using circularity, where values closer to 1 indicate more circular particle shapes. In addition, particle width, height, perimeter, trajectory length, and optical properties derived from RGB intensity values were recorded for each particle.

## Assessment

### Method validation using reference particles (Stokes’ law)

To validate the accuracy of the method, sinking velocities of spherical reference particles with known physical properties were measured and compared to theoretical predictions based on Stokes’ law. Polystyrene beads (size 200-300 µm; density 1.05 g cm^-3^) were used as reference particles, as their sinking behavior in water can be described using physical principles. According to Stokes’ law, the theoretical sinking velocity of a spherical particle in a fluid is given by:

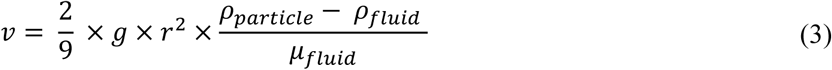

Where v is the terminal sinking velocity, g is the gravitational acceleration (9.81 m s^-2^), r is the particle radius, ρ_particle_ is the particle density, ρ_fluid_ is the fluid density, and μ is the dynamic viscosity of the fluid, which depends on temperature and salinity. This equation assumes low Reynolds numbers (< ∼0.1-0.5, McNown and Malaika 1950), under which viscous forces dominate over inertial forces and fluid motion remains laminar, allowing particles to rapidly reach a constant terminal velocity. These conditions are generally fulfilled for small particles in calm water.

To assess the accuracy of our measurement system, the PSSC was filled with Milli-Q water maintained at 22°C using the integrated temperature control system. Reference particles were introduced from the top of the chamber, and their sinking trajectories were recorded using the tracking and post-processing workflow described above.

The measured and for wall effects corrected velocities of sinking beads were then compared to the theoretically calculated Stokes’ law range of 1.19-2.68 mm s^-1^. Only trajectories longer than 2.5 mm were included in the comparison to reduce uncertainty from short particle tracks. Individual bead trajectories showed some variability, but the central distribution of measured velocities closely matched the theoretical range (Figure 3). The interquartile range of measured velocities (1.76–2.67 mm s⁻¹) fell entirely within the theoretically predicted range, and the median measured velocity was 2.24 mm s^-1^. This agreement indicates that the setup and script reliably capture particle sinking behavior under controlled conditions.

**Figure 3:**
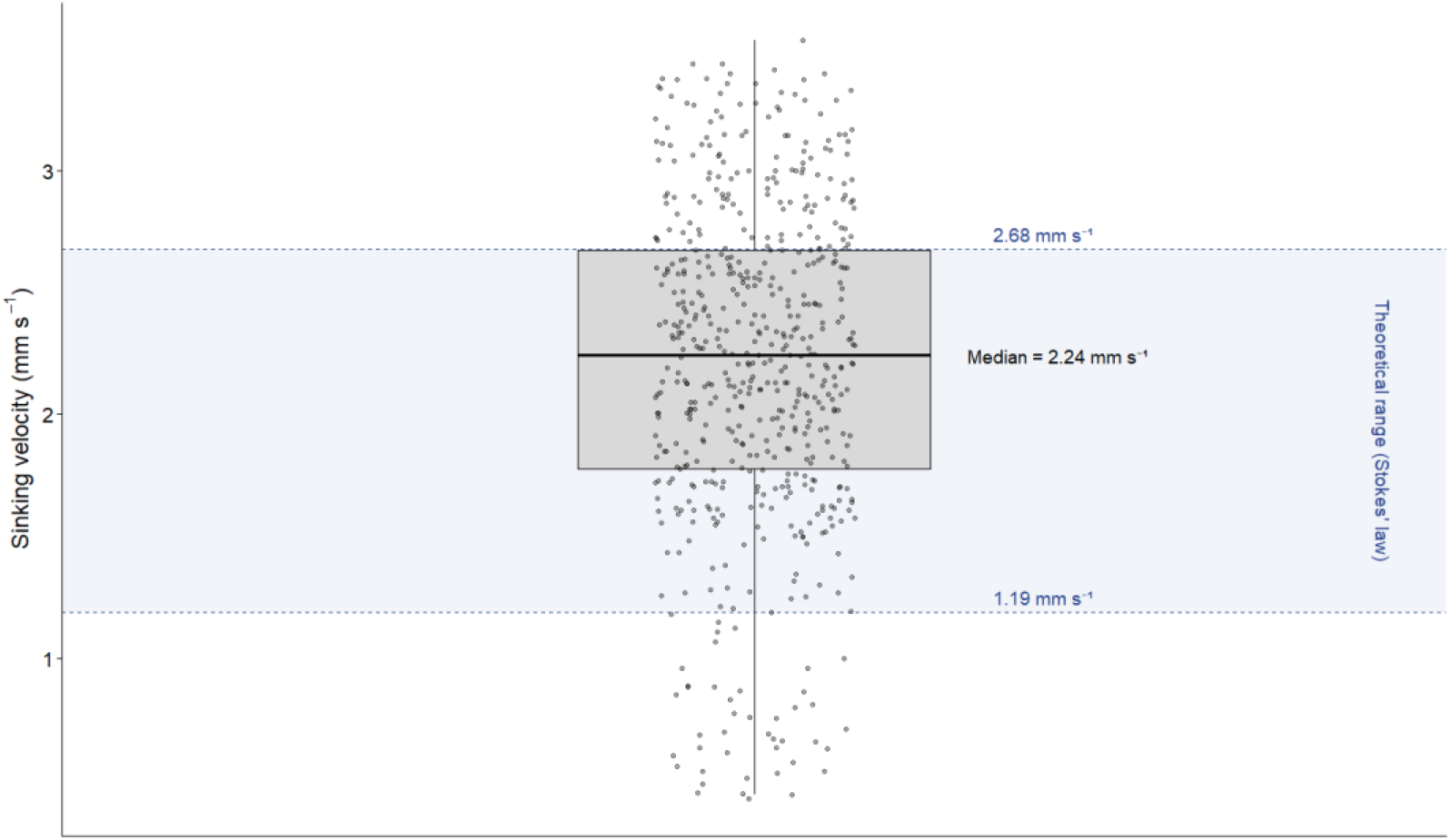
Sinking velocities of polystyrene reference beads measured in the particle sinking speed chamber (PSSC). The shaded area and dashed lines show the theoretical Stokes-law velocity range calculated for the bead size range of 200 – 300 µm. The points represent individual bead trajectories longer than 2.5 mm. The box shows the interquartile range, with the horizontal line indicating the median velocity. The interquartile range of measured velocities fell within the theoretical Stokes-law prediction range, supporting the accuracy of the sinking velocity measurements.

Minor deviations between the measured and theoretical velocities may be due to slight deviations from an ideal spherical shape, still minor wall effects within the PSSC, and uncertainties in assumed particle density values, particle size, and fluid properties. Nevertheless, the overall correspondence supports the reliability of the method for quantifying sinking velocities of small particles.

### Field application

The applicability of the method under field conditions was tested during research cruises and coastal deployments in the Baltic Sea. The settling tank system was successfully used on a research vessel (R/V Aranda) and on small boats in coastal environments. In the field, water was filled into the tanks running the pump with a handheld aggregate. Because smaller boats caused more movements due to waves, we started the 2 h settling period only after returning to the harbor. This minimized water movement inside the tanks and ensured reliable particle settling. During filling and transporting in the field, the tanks remained stable. It was easy to handle and did not need any special infrastructure.

As an example of successful fractionation under field conditions, the chlorophyll a concentration consistently differed between suspended and fast-sinking fractions at both sampling depths and across all sampling dates (Figure 4). Its concentration in the fast-sinking fractions was on average 62% higher at 5 m and 88% higher at 25 m compared to the suspended fraction. The consistent enrichment pattern indicates effective particle separation and suitability for use on small platforms.

**Figure 4:**
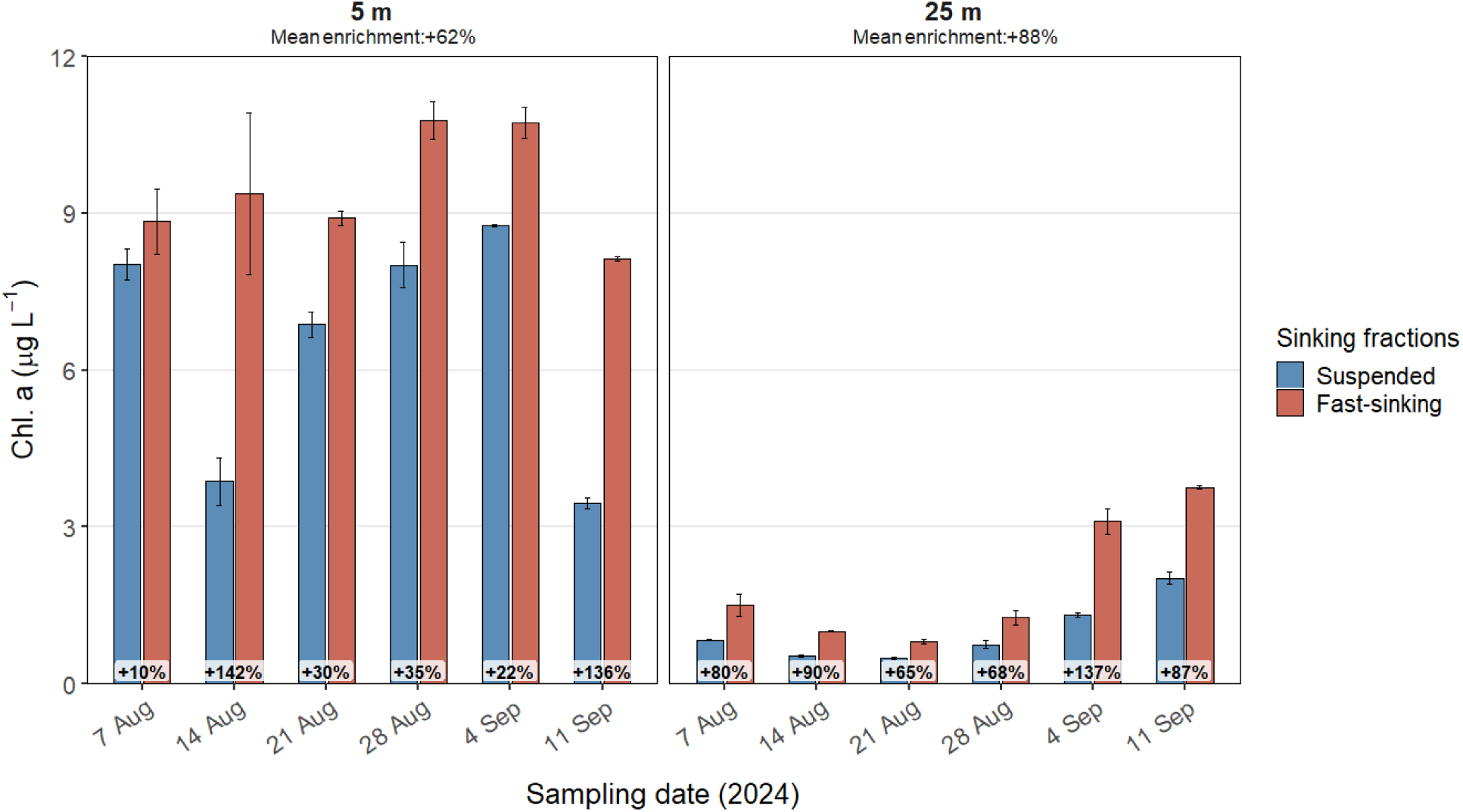
Chlorophyll a concentrations in suspended (blue bars) and fast-sinking (red bars) particle fractions collected with settling tanks during coastal field deployments in Tvärminne (Baltic Sea) in 2024. Measurements are shown for 5 m and 25 m water depth. Error bars indicate standard deviation from triplicate measurements. Percentage values indicate the enrichment of chlorophyll a in the fast-sinking fraction relative to the suspended fraction for each sampling date. Mean enrichment across all sampling dates is given for each depth. The consistent enrichment of chlorophyll a in the fast-sinking fraction demonstrates successful particle fractionation under field conditions.

The imaging system for sinking velocity measurements was tested both on research vessels and in the lab. During research cruises in 2025, particle trajectories did not show visible effects of the vessel movement. This stability was likely supported by closing the PSSC system without air bubbles, which minimized internal water movement and reduced the influence of external disturbances. However, during a later cruise in 2026, some trajectories showed still movement patterns that may have been influenced by vessel vibrations, indicating that measurement stability can depend on sea conditions and platform movement. On smaller boats, stronger vessel movements limit direct measurements onboard. In these cases, samples for sinking velocity measurements were collected from settling tanks in the harbor and then measured under stable land conditions.

### Overcoming hydrodynamic artefacts (temperature control)

During our measurements with this setup, a certain proportion of tracked particles showed upward moving trajectories, indicating convective motion within the system. This pattern can arise from temperature differences between the water sample and the surrounding conditions, which create density gradients and induce convective circulation within the PSSC. As a result, particle movement is not only influenced by gravitational settling but also by vertical fluid motion.

To reduce these effects, measurements were conducted in a climate-controlled room set to in situ water temperatures. Sampled water was stored under the same conditions prior to analysis. These measures reduced the occurrence of visible advection, but even small thermal gradients can affect particle behavior, so temperature control is critical. The integrated cooling system stabilized the temperature conditions during the particle tracking and reduced density differences within the chamber, minimizing advection. Most particles had consistent downward trajectories, enabling a more reliable estimation of actual sinking velocities. Because the samples were not fixed, some trajectories reflected active movement by organisms. Such trajectories were excluded during post-processing using the predefined filtering criteria.

For stable measurements, the cooling system should be also operated when sample temperatures are close to room temperature. Even small temperature differences between sample, chamber walls, and surrounding air can generate thermal gradients that affect particle motion. We also noted that cooling under humid conditions can lead to condensation on the front wall of the PSSC interfering with the optical measurements. The risk of condensation increases with larger temperature differences between the sample and surrounding air and with increasing humidity. Therefore, measurements benefit from keeping room and sample temperature as similar as possible, particularly when working with cold water samples. Under humid conditions, a dehumidifier could also reduce the risk of condensation.

### Output of imaging workflow

The imaging workflow made it possible to simultaneously track and characterize large numbers of sinking particles. From the extracted trajectories, we determined sinking velocities, particle sizes, circularities for each particle. Additional RGB-based information was also recorded. Particle velocities and properties obtained from multiple field measurements covered a broad range (Figure 5). Sinking velocities ranged from 0.01 to 290 m d^-1^, with 99% of particles with velocities between 0.24 and 42.89 m d^-1^. Particle sizes spanned more than two orders of magnitude, reaching up to 660 µm ABD, while 99% of particles were smaller than 147 µm. Circularity values ranged up to 0.95, with 99% of particles exhibiting circularities between 0.32 and 0.94. These distributions show that the workflow captures substantial variability in particle sinking behavior and morphology across natural samples. An example output file generated from a single field measurement in the Baltic Sea is provided in the Supporting Information (Table S1).

**Figure 5:**
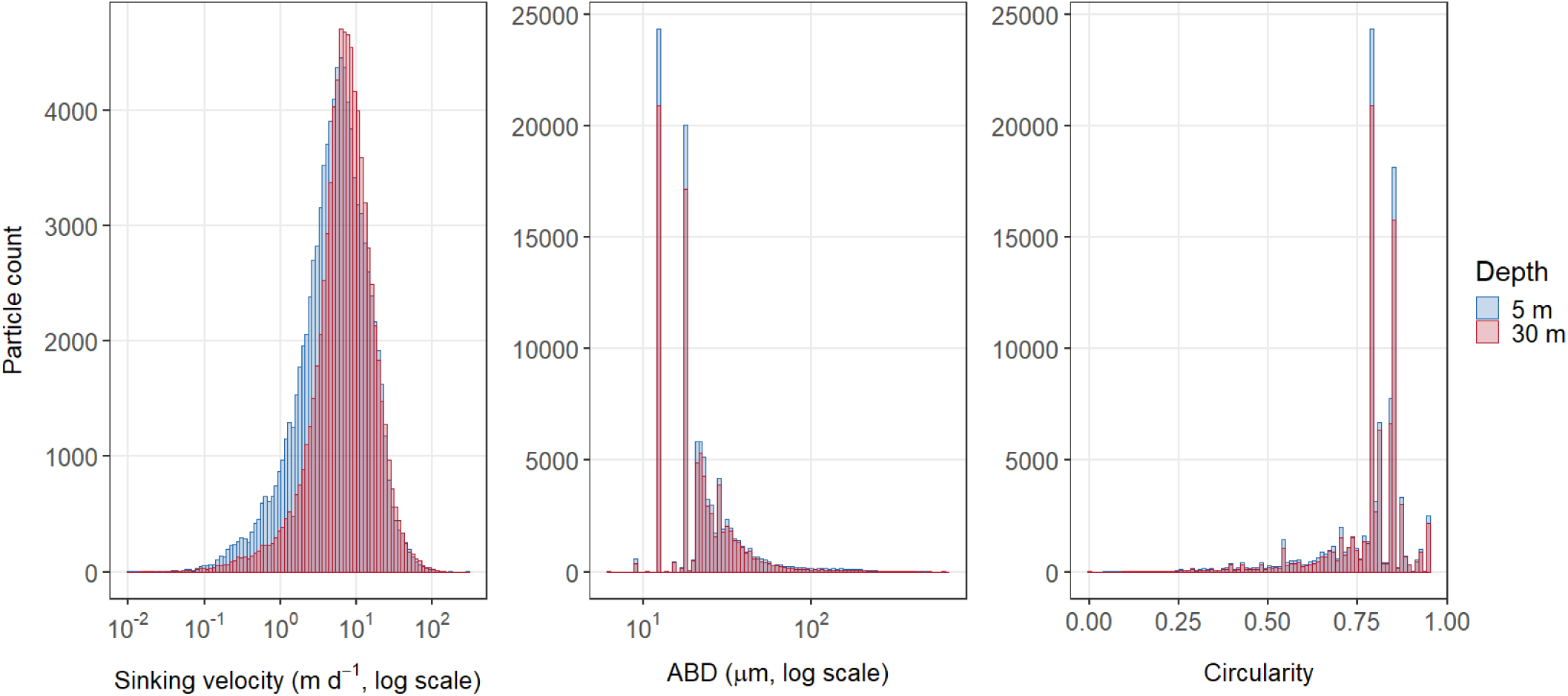
Distributions of sinking velocities, area-based diameter (ABD), and circularity for sinking particles measured at 5 m (blue) and 30 m (red) depth. Sinking velocity and ABD are shown on logarithmic x-axes. The broad distributions show the range of particle velocities and properties that can be measured simultaneously with the imaging workflow.

### RGB-based particle properties

In addition to sinking velocity, size, and shape, the script extracts RGB intensity information from each detected particle. In principle, these data could provide information on particle color and transparency characteristics and may therefore offer additional insights into particle composition. However, meaningful comparison of RGB values between measurements requires highly standardized imaging conditions, including consistent lighting and background characteristics. Small variations in these conditions can influence the recorded RGB values and limit comparability among videos. The setup used in the present study was not optimized for comparison of RGB values among measurements. Consequently, RGB-based particle properties were not evaluated further in the present study. Nevertheless, these variables were retained in the script. Future implementations could readily achieve the required standardization through calibrated light systems and standardized backgrounds, potentially enabling additional optical particle characterization.

### Strengths, recommendations, and limitations

The major advantage of our design for investigating sinking particle dynamics lies in the low-cost and lightweight setup, making it easy to transport and operate with limited amount of infrastructure. An additional strength is the temperature control around the PSSC reducing the problem of small-scale advection that affects the measurements.

Detailed assembly instructions and data processing workflows, including the particle-tracking and post-processing script, are provided to enable consistent implementation and reproducibility. Because of its modular design, the setup can be easily adapted to fit the needs of individual experiments and leaves room for further improvements, such as using a different camera depending on the spatial resolution needed. Sinking velocities can be determined for a large number of particles per measurement, while particle properties such as size and shape, as well as additional RGB-based information, are extracted simultaneously. This enables a comprehensive and efficient characterization of many particles and their sinking behavior. The smallest particles that can be reliably detected depend on image quality, particle contrast, and the chosen camera and image-processing settings. As a result, the effective lower size limit may differ between experimental setups.

When applying the method, some methodological considerations and sources of uncertainty should be considered. Exact determination of sinking velocities depends on precise calibration of the imaging system, especially the distance between camera and PSSC and any zooming of the camera affecting the length of the field of view. That directly affects the conversion from pixels to µm. Small calibration errors can alter the calculated velocities. In addition, comparatively low particle densities and relatively large chamber dimensions should be considered. These conditions help to minimize hydrodynamic interactions between nearby particles, which can influence sinking velocities at high particle concentrations (Happel and Brenner 1983; Bach et al. 2012). Still, a sufficiently large number of particles should be measured to account for variability in particle orientation during sinking (Bach et al. 2012).

Another source of uncertainty is that non-spherical particles can sink differently depending on their orientation in the water column. For example, elongated particles such as fecal pellets may sink faster when vertically oriented than when horizontally oriented (Holland 2010). Because particles are tracked only over relatively short distances, such orientation effects can increase the variability in measured sinking velocities. Image disturbances such as background noise caused by chamber imperfections or construction materials (e.g. glue residues) can also affect particle detection. However, these effects can largely be reduced during the video processing step. In addition, the smallest particles that can be detected depend on image quality, particle contrast, camera settings, and image-processing parameters. Therefore, the effective lower size limit may vary between applications and should be assessed for each experimental setup.

Possible artifacts of the method may arise from the sampling design and tracking approach. As sinking particles may move in and out of the camera focus, tracking artefacts may occur. This can potentially lead to fragmented trajectories and an overestimation of the particle numbers. It is possible to make particle tracking more conservative in the script, e.g. only counting the full-length tracks (whole field of view), but that would likely underestimate the total sinking particles. This is why this method is not optimized for determining absolute particle abundances, although the overall order of magnitude of particle concentrations is still captured.

One limitation of the sampling method is that it may affect particle integrity. Although the peristaltic pump used here handles the material relatively gentle, very fragile or large aggregates may partly break apart during transfer into the settling tanks or when transferred to the PSSC. This should be kept in mind when interpreting particle size distribution and flux behavior. The method is primarily suited for coastal and shelf environments where it is mainly primary particles and is less applicable to deep ocean sampling where there are more fragile aggregates. This is also because it relies on pumping water to the surface which becomes increasingly challenging with depth. In our example, we let the pump run for 2 min to ensure we get water from 30 m depth. Increasing the depth would increase this pumping time, which could influence particle properties.

Some limitations to our method also arise from the analytical design of the approach. Chemical properties of individual particles cannot be resolved. Therefore, biogeochemical information is limited to bulk analyses of the particle fractions from the settling tank. However, bulk measurements still provide complementary insights and are sufficient for many applications related to carbon export.

## Discussion

In this study, we present and validate a low-cost, modular system for separating sinking particle fractions and measuring individual particle sinking velocities under controlled conditions. The separation of field-derived particle fractions provides access to natural particle assemblages and enables associated biogeochemical bulk analyses. Optical tracking then allows the determination of sinking velocities in relation to particle-specific properties. Together, this approach links physical particle dynamics with biogeochemical characteristics and provides a practical tool to investigate processes relevant to the BCP under controlled but realistic conditions. The primary advance of the presented method is not the measurement of sinking velocities itself, but the reduction of logistical and financial barriers that currently limit particle-resolved observations of carbon export dynamics in coastal systems.

Existing approaches for measuring particle sinking velocities involve trade-offs between ecological realism, information content, logistical requirements, and cost. The method presented here is composed of two parts, first the settling tank to concentrate fast sinking particles and second the sinking speed measurements. The settling tank is comparable to more established approaches such as the marine snow catcher (Lampitt et al. 1993; Romanelli et al. 2023). Similar to marine snow catchers, the settling tank collects natural particle assemblages directly from the target depth and allows subsequent biogeochemical characterization of separated particle fractions. The marine snow catcher was developed to collect intact marine aggregates with minimal hydrodynamic disturbance. They are typically similar in size to our tanks (∼110 L) and collect the water directly from the desired depth. However, they are relatively expensive (∼100-fold compared to our tanks) and require heavy lifting capabilities (crane and winch), which limit their application in shallow coastal systems. In contrast, our settling tank approach uses inexpensive components and can be deployed from small boats without specialized infrastructure.

The sinking speed measurements in a dedicated PSSC with a surrounding water TC chamber complement existing methods for determining sinking velocities or particles/aggregates. Similar to other ex situ approaches, such as sedimentation columns or vertical flow systems, the system provides controlled measurement conditions and links sinking velocities to particle-specific properties. The low cost of our system is comparable to the traditional picking of particles/aggregates followed by manual measurement of sinking speed over a predefined distance. However, our approach has the advantage of being able to measure the sinking speeds of large numbers of particles per measurement (typically several thousand particles, up to more than 18,000 particles in the present study). Compared to other imaging devices such as the FlowCam which has successfully been used for the same purpose (Bach et al. 2012), our system can be obtained at a fraction of the price (<1%) and with the added benefit of having the temperature control surrounding the PSSC minimizing particle advection during measurements.

Our method can be combined with additional analysis, giving potential for new insights into particle dynamics and carbon cycling beyond sinking speed alone. The separated fractions can be used to analyze elemental stoichiometry and respiration rates to assess the particle composition and the balance between carbon export efficiency and remineralization. Biological analyses could also be added to link sinking properties and particle characteristics to biological communities and processes. These could include DNA- or RNA-based sequencing methods to determine phytoplankton/bacteria community composition and active gene regulation. Furthermore, the role of ballasting minerals and substances such as transparent exopolymer particles (TEP) could be investigated, as minerals enhance particle density, whereas TEP promotes aggregation and particle growth (Alldredge et al. 1993; Lombard et al. 2013).

In conclusion, the presented method is low-cost, modular, and lightweight, and can be assembled from widely available materials. It can be easily transported and deployed from small platforms without the need for heavy equipment or large water sampling systems. This reduces logistical and financial barriers that can limit the application of existing methods, particularly in resource-limited research settings. The successful application under field conditions in coastal environments has demonstrated that particle sinking dynamics can be measured efficiently from small vessels or boats. That makes the method ideal for shallow and undersampled regions. Expanding these observations across diverse coastal systems will help to close existing gaps in our knowledge and lead to a more complete picture of carbon export efficiency in these highly variable environments.

## Supporting information

Script S1

## Acknowledgements

We would like to thank E. Lundt for his valuable help in developing the script, as well as for many insightful discussions and ideas. We are also grateful to A. Fernández-Carrera for her thoughtful feedback and support in reviewing and improving the manuscript. In addition, we thank the staff at the Tvärminne Zoological Station for their assistance in the field and with sample collection. This study was financed by the Research Council of Finland (decision 354272) and used infrastructure part of the FINMARI research infrastructure consortium.

Chat GPT was used to help formulate part of the text.

