## Supplementary Information for "A low-cost method for collecting and measuring sinking speed of particles in coastal zones"

### *Figures*

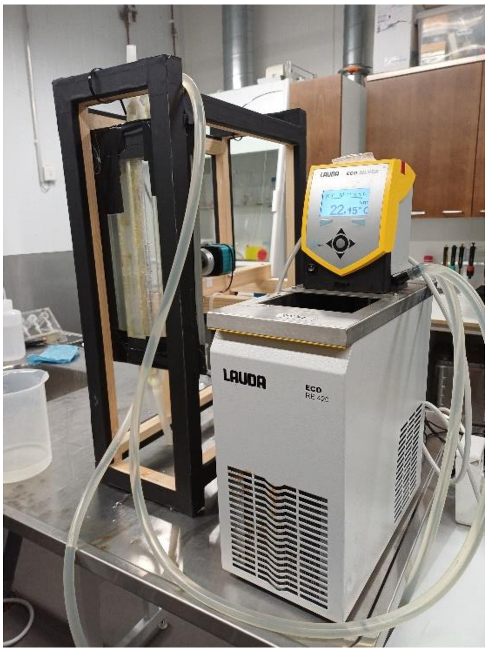


Figure S1: Additional photograph of the sinking velocity measurement setup, including the polycarbonate boards forming the SSC and surrounding TC chamber, the supporting wooden frame, camera and the circulating water temperature control unit (LAUDA RE 420 SN) used for temperature stabilization during measurements.

### *Supporting files*

Script S1. Python script used for automated particle detection, trajectory tracking, and calculation of sinking velocities and particle properties from video recordings.

Table S1. Example output file produced by the particle-tracking workflow from a measurement of the fast-sinking particle fraction collected in the Baltic Sea (station US5B; 62.58615° N, 19.96883° E) from 5 m depth on 3 June 2025. The file contains the particle properties and sinking velocities calculated with the supplied python script.
