## Supplementary material for "A low-cost method for collecting and measuring sinking speed of particles in coastal zones": Script S1

"""
Supplementary Script S1

Particle tracking and characterization of sinking particles from video recordings.


Description
-----------
This script detects, tracks, and characterizes individual particles in video recordings. Tracked particles are used to calculate sinking velocities and particle properties, including size, shape, and color-based descriptors.

Input
-----
- Video files (.mov)


Output
------
- CSV files containing particle trajectory data and derived particle properties including sinking velocity, area-based diameter (ABD), equivalent spherical diameter (ESD), circularity, and particle color properties.


Output units
------------
If a meter-per-pixel ratio is provided, distances are exported in meter and velocities in m/s and m/d. If no ratio is provided, pixel-based values are exported.


Output variables
----------------
dx, dy                      Horizontal and vertical displacement
sinking                     Particle classified as sinking based on vertical and horizontal displacement criteria
width, height               Mean particle dimensions
lifetime_frames             Tracking duration (frames)
lifetime_seconds            Tracking duration (seconds)
average_y_speed             Vertical velocity
average_y_speed_m/d         Vertical velocity (m/d)
wall_corrected_y_velocity   wall-corrected velocity
path_length                 Total particle path length
perimeter                   Particle perimeter
circularity                 Particle circularity
ESD                         Mean Feret diameter
ABD                         Area-based diameter
mean_color_bgr              Mean particle color
color_covariance            Color variability descriptor


Requirements
------------
Python 3.x
OpenCV
NumPy
Pandas
 

Additional packages
-------------------
pyarrow (optional, for parquet export)


Associated manuscript
---------------------
Manuscript in preparation:

Lemke, J. & Spilling, K.
"A low-cost method for collecting and measuring sinking speed of particles in coastal zones."


Authors
-------
Josephin Lemke
Evert Lundt


Version
-------
Version 1.0
Date: June 2026
"""


# ============================================================
# Imports
# ============================================================

import cv2
import glob
import math
import numpy as np
import pandas as pd
from typing import List, Optional, Tuple, Union


# ============================================================
# Helper functions
# ============================================================

def encode_mask(mask: np.ndarray) -> Tuple[int, int, bytes]:
    """
    Convert a binary mask (H×W, uint8 0/255 or 0/1) to a compact bytes representation.
    Returns: (height, width, packed_bytes)
    """
    h, w = mask.shape
    flat = (mask > 0).astype(np.uint8).flatten()
    packed = np.packbits(flat).tobytes()
    return h, w, packed


# ============================================================
# Particle class and tracking information
# ============================================================

class Blob:
    def __init__(self, x: int = 0, y: int = 0, width: int = 0, height: int = 0, first_frame: int = 0):
        self.x = int(x)
        self.y = int(y)
        self.width = int(width)
        self.height = int(height)

        # Tracking meta
        self.first_frame: int = int(first_frame)
        self.last_frame: int = int(first_frame)
        self.confidence: int = 10  # grows on match, decays when missed

        # Path tracking
        self.x_start, self.y_start = self.center()
        self.x_history, self.y_history = self.center()
        self.path_length: float = 0.0
        self.path_count: int = 10  # update path every 10 frames to suppress noise

        # Running averages for size
        self.width_sum: float = float(self.width)
        self.height_sum: float = float(self.height)

        # Shape/color descriptors (set lazily)
        self.shape: Union[int, bytes] = -1  # -1 means not yet set; else bytes from encode_mask
        self.sw: int = -1
        self.sh: int = -1
        self.mu: Optional[np.ndarray] = None          # shape (3,)
        self.Sigma: Optional[np.ndarray] = None       # shape (3,3)

    def __repr__(self):
        return (f"Blob(x={self.x}, y={self.y}, w={self.width}, h={self.height}, "
                f"first={self.first_frame}, last={self.last_frame}, conf={self.confidence})")

    def right(self) -> int:
        return self.x + self.width

    def center(self) -> Tuple[float, float]:
        return (self.x + self.width / 2.0, self.y + self.height / 2.0)

    def nearest(self, bboxes: List[Tuple[int, int, int, int]], max_distance: float) -> Optional[Tuple[int, int, int, int]]:
        """
        Return the closest bbox within max_distance (in pixels), or None.
        """
        bestd = max_distance ** 2
        bestmatch = None
        ox, oy = self.center()
        for box in bboxes:
            x, y, w, h = box
            nx = x + w // 2
            ny = y + h // 2
            dist = (nx - ox) ** 2 + (ny - oy) ** 2
            if dist < bestd:
                bestmatch = box
                bestd = dist
        return bestmatch

    def _compute_roi_stats_once(self, frame: np.ndarray, mask: np.ndarray):
        """
        Compute color mean/covariance and encoded shape on the ROI. Called once per blob.
        Guarded to avoid NaNs/empty ROI.
        """
        # Defensive clipping just in case
        h_img, w_img = mask.shape
        x0, y0 = max(self.x, 0), max(self.y, 0)
        x1, y1 = min(self.x + self.width, w_img), min(self.y + self.height, h_img)
        if x1 <= x0 or y1 <= y0:
            return

        roi_mask = mask[y0:y1, x0:x1]
        roi_frame = frame[y0:y1, x0:x1]

        # Collect pixels inside mask
        X = roi_frame[roi_mask != 0]
        if X.size == 0:
            return

        # Color mean
        self.mu = X.mean(axis=0)

        # Covariance (guard for few pixels)
        if X.shape[0] >= 2:
            self.Sigma = np.cov(X, rowvar=False, bias=False)
        else:
            self.Sigma = np.zeros((3, 3), dtype=float)

        # Encoded shape
        self.sh, self.sw, self.shape = encode_mask(roi_mask)

    def update(self, box: Tuple[int, int, int, int], frame: np.ndarray, fg_mask: np.ndarray, frame_counter: int):
        """
        Update blob kinematics and (periodically) path length; compute ROI stats once per blob.
        """
        frames_since_detection = frame_counter - self.last_frame 

        x, y, w, h = box
        self.x = int(x)
        self.y = int(y)
        self.width = int(w)
        self.height = int(h)
        self.last_frame = int(frame_counter)

        # Confidence increases on successful association
        self.confidence = min(self.confidence + 6, 100)

        # Keep running sums for average width/height
        self.height_sum += float(self.height)*frames_since_detection
        self.width_sum += float(self.width)*frames_since_detection

        # Update path every N frames to avoid noisy path length
        if self.path_count < 1:
            bx, by = self.center()
            self.path_length += math.hypot(bx - self.x_history, by - self.y_history)
            self.x_history, self.y_history = bx, by
            self.path_count = 10
        else:
            self.path_count -= frames_since_detection

        # Compute color/shape descriptors once (first time we see a stable ROI)
        if isinstance(self.shape, int):  # not yet computed
            # Option A: require at least 10 frames since first_frame for stability
            if (self.last_frame - self.first_frame) >= 10:
                self._compute_roi_stats_once(frame, fg_mask)

    def save(self, collector: Optional[list], fps: float):
        """
        Save blob metrics if it lived long enough. If collector is None, just print.
        Converts numpy arrays to serializable types before saving.
        """
        lifetime_frames = self.last_frame - self.first_frame
        lifetime_seconds = lifetime_frames / fps if fps > 0 else lifetime_frames / 60.0

        # Add the final path segment from last history point to current center
        cx, cy = self.center()
        self.path_length += math.hypot(cx - self.x_history, cy - self.y_history)

        # Avoid division by zero
        avg_w = self.width_sum / max(lifetime_frames, 1)
        avg_h = self.height_sum / max(lifetime_frames, 1)
        # Calculate mean vertical velocity from net vertical displacement 
        # between the first and last detected particle center.
        average_speed = (cy - self.y_start) / max(lifetime_frames, 1)  # px/frame

        record = {
            "dx": cx - self.x_start,
            "dy": cy - self.y_start,
            "width": avg_w,
            "height": avg_h,
            "lifetime_frames": lifetime_frames,
            "lifetime_seconds": lifetime_seconds,
            "average_y_speed_px_per_frame": average_speed,
            "average_y_speed_px_per_second": average_speed *fps if fps > 0 else 60,
            "path_length_px": self.path_length,
            "first_frame": self.first_frame,
            # Serialize arrays
            "mean_color_bgr": self.mu.tolist() if isinstance(self.mu, np.ndarray) else None,
            "color_covariance": self.Sigma.tolist() if isinstance(self.Sigma, np.ndarray) else None,
            "shape_height": self.sh if isinstance(self.sh, int) else None,
            "shape_width": self.sw if isinstance(self.sw, int) else None,
            "encoded_shape": self.shape if isinstance(self.shape, (bytes, bytearray)) else None,
        }

        if collector is None:
            print("Blob:", record)
        else:
            collector.append(record)


# ============================================================
# Video processing and particle detection
# ============================================================

def draw(frame: np.ndarray, fg_mask: np.ndarray, frame_counter: int,
         blobs: List[Blob], bboxes: List[Tuple[int, int, int, int]]):
    """
    Overlay visualization of blobs, centers, trajectories, and mask gradient.
    """
    # Morphological gradient highlights edges cleanly
    kernel = cv2.getStructuringElement(cv2.MORPH_ELLIPSE, (4, 4))
    gradient = cv2.morphologyEx(fg_mask, cv2.MORPH_GRADIENT, kernel)

    # Draw blobs and their path (center → last center)
    for blob in blobs:
        color = (0, 0, 255) if (blob.last_frame - blob.first_frame) < 60 else (255, 0, 0)
        # Rectangle
        cv2.rectangle(frame,
                      (int(blob.x) - 1, int(blob.y) - 1),
                      (int(blob.x + blob.width) + 1, int(blob.y + blob.height) + 1),
                      color, 2)
        # Trajectory segment (from previous center to current center)
        cx, cy = blob.center()
        cv2.line(frame,
                 (int(blob.x_history), int(blob.y_history)),
                 (int(cx), int(cy)),
                 (0, 255, 0), 2)

    # Show frame index
    cv2.putText(frame, str(frame_counter), (10, 30), cv2.FONT_HERSHEY_SIMPLEX, 0.5,
                (255, 255, 255), 1, cv2.LINE_AA)

    # Overlay gradient in red
    frame[gradient == 255] = (0, 0, 255)

    cv2.imshow('Original Frame with Changes', frame)


def process_video(
    video_path: str,
    save_results: bool = True,
    display: bool = False,
    max_distance: float = 20.0,
    var_threshold: float = 10,
    morph_kernel_size: int = 3,
    min_lifetime_seconds: float = 1.0,
) -> None:
    """
    Process a video file, track blobs, and optionally save results to parquet.
    """
    cap = cv2.VideoCapture(video_path)
    ret, frame = cap.read()
    if not ret or frame is None:
        cap.release()
        if display:
            cv2.destroyAllWindows()
        print(f"Could not read first frame: {video_path}")
        return

    fps = cap.get(cv2.CAP_PROP_FPS) or 60.0  # fallback if FPS not available
    min_lifetime_frames = int(round(min_lifetime_seconds * fps))

    bg_subtractor = cv2.createBackgroundSubtractorMOG2(
        history=500, varThreshold=var_threshold, detectShadows=False
    )
    kernel = cv2.getStructuringElement(cv2.MORPH_ELLIPSE, (morph_kernel_size, morph_kernel_size))

    frame_counter = 0
    blobs: List[Blob] = []
    results: Optional[list] = [] if save_results else None

    try:
        while ret:
            # Detect moving particles using background subtraction.
            fg_mask = bg_subtractor.apply(frame, learningRate=0.0001)
            # Clean mask
            fg_mask = cv2.morphologyEx(fg_mask, cv2.MORPH_CLOSE, kernel)
            fg_mask = cv2.morphologyEx(fg_mask, cv2.MORPH_OPEN, kernel)

            contours, _ = cv2.findContours(fg_mask, cv2.RETR_EXTERNAL, cv2.CHAIN_APPROX_SIMPLE)
            bboxes = [cv2.boundingRect(cnt) for cnt in contours]

            # Associate existing blobs
            for blob in blobs[:]:  # iterate over a shallow copy to allow removals
                box = blob.nearest(bboxes, max_distance=max_distance)
                if box is not None:
                    blob.update(box, frame, fg_mask, frame_counter)
                    bboxes.remove(box)
                else:
                    # No match this frame → decay confidence
                    blob.confidence -= 5
                    if blob.confidence <= 0:
                        # Only save if lifetime frames exceed threshold
                        if (blob.last_frame - blob.first_frame) >= min_lifetime_frames:
                            blob.save(results, fps=fps)
                        blobs.remove(blob)

            # Initialize new blobs for unmatched boxes
            for box in bboxes:
                x, y, w, h = box
                blobs.append(Blob(x, y, w, h, first_frame=frame_counter))

            if display:
                draw(frame.copy(), fg_mask, frame_counter, blobs, bboxes)
                if cv2.waitKey(30) & 0xFF == 27:  # ESC to stop
                    break

            # Progress log every 600 frames (~10s @60fps)
            if frame_counter % 600 == 0 and frame_counter > 0:
                print(f"Processed {frame_counter} frames ({frame_counter / fps:.1f} s)")

            # Next frame
            frame_counter += 1
            ret, frame = cap.read()

        # Finalize any remaining blobs when video ends
        for blob in blobs:
            if (blob.last_frame - blob.first_frame) >= min_lifetime_frames:
                blob.save(results, fps=fps)

        # Save results
        if results is not None:
            df = pd.DataFrame(results)
            # Parquet may need pyarrow; if missing, fall back to CSV
            save_csv(df, video_path + "c.txt")
            try:
                df.to_parquet(video_path + "c.parquet")
            except Exception as e:
                print(f"Parquet export failed ({e}); writing CSV instead")
                df.to_csv(video_path + "c.csv", index=False)

        print(f"Done processing {video_path}")

    except Exception as e:
        print(f"Error processing {video_path}: {e}")
    finally:
        cap.release()
        if display:
            cv2.destroyAllWindows()


# ============================================================
# Particle property calculations and data export
# ============================================================

def extract_mask(row):
    w, h = row["shape_width"], row["shape_height"]
    packed = row["encoded_shape"]
    flat = np.unpackbits(np.frombuffer(packed,dtype=np.uint8))[:h*w]
    mask = flat.reshape((h, w))
    mask = np.pad(mask, ((1, 1), (1, 1)))
    return mask


def feret(cnt):
    cnt = cnt.astype(np.float32)

    # === Compute Feret diameters at 36 angles ===
    angles_deg = np.arange(0, 180, 5)
    feret_diameters = []

    for angle in angles_deg:
        theta = np.deg2rad(angle)
        # Direction vector
        direction = np.array([math.cos(theta), math.sin(theta)], dtype=np.float32)

        # Project contour points onto direction vector
        proj = cnt @ direction

        # Feret = max projection - min projection
        feret = proj.max() - proj.min()
        feret_diameters.append(feret)
    return sum(feret_diameters)/len(feret_diameters)


def save_csv(df, file):
    global ratio, wall_distance  # ratio = meters per pixel, wall_distance = R
    result = []
    print(ratio, wall_distance)
    try:
        for row in df.iterrows():
            row = row[1]
            mask = extract_mask(row)
            cnt, _ = cv2.findContours(mask, cv2.RETR_EXTERNAL, cv2.CHAIN_APPROX_SIMPLE)
            if not cnt:
                continue

            c = cnt[0]

            perimeter_px = cv2.arcLength(c, True)
            contour_area_px = cv2.contourArea(c)
            mean_feret_px = feret(c)
            circularity = (4*math.pi*contour_area_px)/perimeter_px**2 if perimeter_px != 0 else None

            # Compute wall-corrected velocity
            wall_corrected_velocity = None
            if wall_distance is not None:
                # r = radius of equivalent-area circle
                r = math.sqrt(contour_area_px * ratio**2 / math.pi)

                k = 1.004
                R = wall_distance

                denominator = (1 - k * r / R)

                if denominator != 0:
                    wall_corrected_velocity = row["average_y_speed_px_per_second"] * ratio / denominator

            if ratio is not None:
                result.append({
                    "dx": row["dx"] * ratio,
                    "dy": row["dy"] * ratio,
                    # Classify particles as sinking when vertical displacement is positive
                    # and horizontal displacement remains below the defined threshold.
                    "sinking": abs(row["dx"] * ratio) < 0.00020693 and row["dy"] > 0,
                    "width": row["width"] * ratio,
                    "height": row["height"] * ratio,
                    "lifetime_frames": row["lifetime_frames"],
                    "lifetime_seconds": row["lifetime_seconds"],
                    "average_y_speed": row["average_y_speed_px_per_second"] * ratio,
                    "average_y_speed_m/d": row["average_y_speed_px_per_second"] * ratio * 86400,
                    "wall_corrected_y_velocity": wall_corrected_velocity,
                    "wall_corrected_y_velocity_m/d": wall_corrected_velocity * 86400, #86400 seconds in a day
                    "path_length": row["path_length_px"] * ratio,
                    "first_frame": row["first_frame"],
                    "perimeter": perimeter_px * ratio,
                    "circularity": circularity,
              	    # ESD is calculated as the mean Feret diameter.
                    "ESD": mean_feret_px * ratio,
                    # ABD is calculated from projected particle area.
                    "ABD": 2 * math.sqrt((contour_area_px * ratio**2) / math.pi),
                    "mean_color_bgr": row["mean_color_bgr"],
                    "color_covariance": row["color_covariance"]
                })
            else:
                result.append({
                    "dx_px": row["dx"],
                    "dy_px": row["dy"],
                    "width_px": row["width"],
                    "height_px": row["height"],
                    "lifetime_frames": row["lifetime_frames"],
                    "lifetime_seconds": row["lifetime_seconds"],
                    "average_y_speed_px": row["average_y_speed_px_per_second"],
                    "wall_corrected_y_velocity_px": wall_corrected_velocity,
                    "path_length_px": row["path_length_px"],
                    "first_frame": row["first_frame"],
                    "perimeter_px": perimeter_px,
                    "circularity": circularity,
                    "ESD_px": mean_feret_px,
                    "ABD_px": 2 * math.sqrt(contour_area_px / math.pi),
                    "mean_color_bgr": row["mean_color_bgr"],
                    "color_covariance": row["color_covariance"]
                })
        print(file)
        out_df = pd.DataFrame(result)
        out_df.to_csv(file, index=False)

    except Exception as e:
        print(e)


# ============================================================
# User-defined parameters
# ============================================================
#
# DISPLAY:
#     Show particle tracking during processing.
#
# SAVE_RESULTS:
#     Save output files.
#
# ratio:
#     Meter-per-pixel calibration factor.
#
# wall_distance:
#     Distance between particle and chamber wall used for
#     wall-effect correction.
#
# max_distance:
#     Maximum linking distance between detections (pixels).
#
# var_threshold:
#     Sensitivity of the background subtraction algorithm.
#
# morph_kernel_size:
#     Morphological filtering kernel size.
#
# min_lifetime_seconds:
#     Minimum particle tracking duration retained for analysis.
#
# ============================================================

if __name__ == "__main__":
    DISPLAY = False  # toggle visualization
    SAVE_RESULTS = True
    # Use exact file or a glob pattern like '*.mov'
    mov_files = glob.glob('*.mov')
    
    # Spatial calibration factor (m px⁻¹).
    # Example: 0.00000806 for a calibration of 720 pixels = 5.8 mm.
    ratio = 0.00000806
    try:
        ratio = float(ratio)

        # Distance from the particle to the chamber wall (m), assuming the
        # particle is located at the center of the camera field of view.
        # Used for wall-effect correction.
        wall_distance = 0.00875

    except ValueError:
        print("No or invalid ratio. Continuing without conversion")
        ratio = None
        wall_distance = None

    
    if not mov_files:
        print("No video files matched the pattern.")
    else:
        for video_file in mov_files:
            process_video(
                video_file,
                save_results=SAVE_RESULTS,
                display=DISPLAY,
                max_distance=20.0,
                var_threshold=40,
                morph_kernel_size=3,
                min_lifetime_seconds=1.0,
            )
